# Pre-existing cutaneous conditions could increase the risk for SARS-COV-2 infection

**DOI:** 10.1101/2020.06.30.181297

**Authors:** Qiannan Xu, Lihong Chen, Li Zhang, Mengyan Hu, Xiaopan Wang, Qi Yang, Yunchen Le, Feng Xue, Xia Li, Jie Zheng

## Abstract

Since the end of 2019, COVID-19 pandemic caused by the SARS-CoV-2 emerged globally. The angiotensin-converting enzyme 2 (ACE2) on the cell surface is crucial for SARS-COV-2 entering into the cells. We use SARS-COV-2 pseudo virus and humanized ACE2 mice to mimic the possible transmitting of SARS-COV-2 through skin based on the data we found that skin ACE2 level is associated with skin pre-existing cutaneous conditions in human and mouse models and inflammatory skin disorders with barrier dysfunction increased the penetration of topical FITC conjugated spike protein into the skin. Our study indicated the possibility that the pre-existing cutaneous conditions could increase the risk for SARS-COV-2 infection.

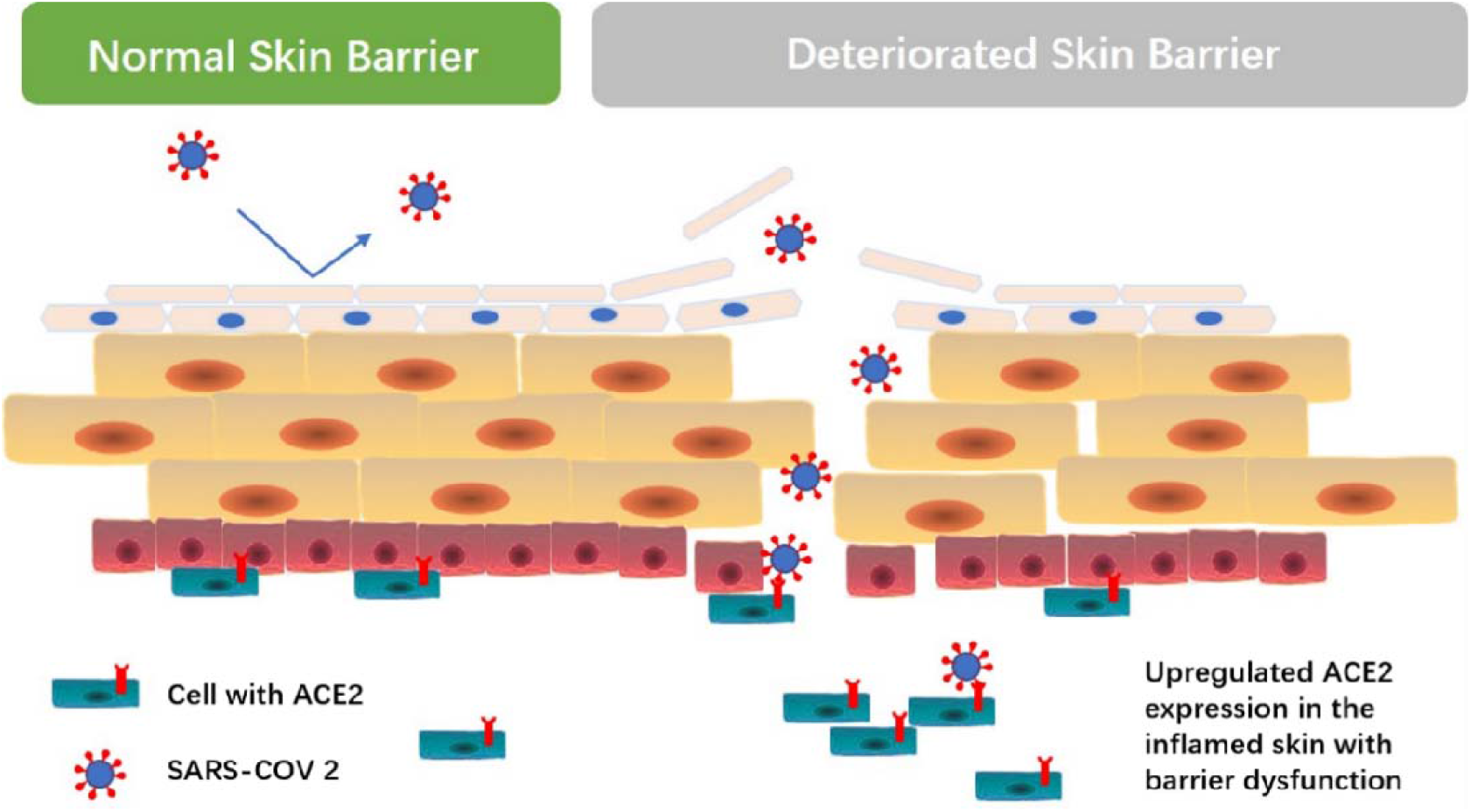

## Introduction

Since the end of 2019, a new corona virus emerged and widely spread throughout the world (Li et al., 2020). This virus was further recognized as SARS-CoV-2 (Wang et al., 2020). With its spike protein (Yan et al., 2020) SARS-CoV-2 could bind angiotensin-converting enzyme 2 (ACE2) on alveolar epithelial type II cells (Ziegler et al., 2020) in the lung and cause severe lung injury, multi-organ failure and death (Huang et al., 2020). Respiratory tract was recognized as the major way of transmission (Chan et al., 2020). Higher expression of ACE2 was thought to be associated with elevated susceptibility to SARS-CoV-2 (Smith et al., 2020). However, wearing a mask only reduce the risk of SARS-CoV-2 through respiratory tract (Javid et al., 2020). The possibility of SARS-CoV-2 infection through other tissue, especially in those with expression of ACE2, could not be excluded (Qi et al., 2020). Thus, either downregulation of ACE2 expression or prevention ACE2 expressing cells from contacting of SARS-CoV-2 could be a solution for preventing the disease (Wu et al., 2020). One of the well observed potential target organ of SARS-CoV-2 is gut (Lamers et al., 2020) because it held a high ACE2 expression and several symptoms related to digestive systems as diarrhea, vomiting, and abdominal pain were observed in COVID-19 patients (Wong et al., 2020). Similar to gut, skin could also express ACE2 (Grzegrzolka et al., 2013; Hamming et al., 2004; Xue et al., 2020) and cases were reported that patients with COVID-19 could have vascular skin symptoms (Bouaziz et al., 2020), varicella-like exanthem (Marzano et al., 2020), and pernio-like skin lesions (Freeman et al., 2020) which were all symptoms occurred on skin. Besides that, SARS-COV-2 particles were found in the cytoplasm of endothelial cells of COVID-19 patients on electron microscopy (Colmenero et al., 2020). And skin rashes could be an orphan symptom in some patient (Tatu et al., 2020). Thus, it indicated that skin as a first line defensing barrier of human body which constantly exposure to various pathogens (Kupper and Fuhlbrigge, 2004) could be another potential target organ of SARS-CoV-2.

When an organ become a potential organ of transmitting SARS-CoV-2 there were to be concern that nowadays widely used biological agents which treating diseases of that organ should elevate the susceptibility to SARS-CoV-2 (Potdar et al., 2020). Because using biologics means more vulnerable to various pathogens (Click and Regueiro, 2019). Psoriasis as an immune-mediated, genetic related disease manifesting in the skin with high prevalence, chronicity, disfiguration, disability, and associated comorbidity (Boehncke and Schön, 2015), biologics played an important role in treating it. However, with the pandemic of COVID-19, whether using biologics could elevated the susceptibility to SARS-CoV-2 remained unknown (Chat et al., 2020). Using biologics treat skin disease in COVID-19 pandemic has its controversial portrait. Although cases reported severe infection occurred after biologics treatment (Bardazzi et al., 2020; Byrd et al., 2018; Lebwohl et al., 2020), biologics could somehow reduce the possibility of transmitting pathogens through skin ? After biologics treatment the inflammation were gone and deteriorated skin barrier function was back to normal. The widened physical gap between keratinocytes of dysfunctional skin barrier returned to the normal skin barrier with tightly conjunct keratinocytes (Dainichi et al., 2018). This contribute to the defensing of various pathogens from skin (Byrd et al., 2018), herein during the COVID-19 pandemic could be SARS-CoV-2.

## Results

### S protein transmitted through the skin of inflammatory skin disorders with barrier dysfunction mouse models

The attachment of spike protein (S protein) with ACE2 is necessary for the transmission of SARS-COV-2. Then could the skin inflammatory disorder with barrier dysfunction affect the S protein entering skin? The influence of skin inflammation and barrier function to the binding of ACE2 and SARS-COV-2 was evaluated. Mouse model of skin inflammatory disorder with barrier dysfunction induced by imiquimod was established and group control was established as applied with vehicle cream (Vaseline Lanette cream, Fargon). Then FITC conjugated S protein was applied to the skin. After that the treated skin was prepared for an immunofluorescence stain. The FITC conjugated S protein was detected depositing in the dermal of mouse model with inflammatory skin disorder and deteriorated skin barrier function (Fig. 1B) while there was none immunofluorescence intensity detected in the mouse model with normal barrier function and without skin inflammatory disorder (Fig. 1A). The FACS scan result of the skin FITC conjugated S protein marked cells were significantly higher in the skin barrier dysfunction group than the control group while by applying the skin barrier recovery moisture the quantity of FITC conjugated S protein permeated the skin barrier significantly reduced (Fig. 4, C to F).

**Fig. 1.**
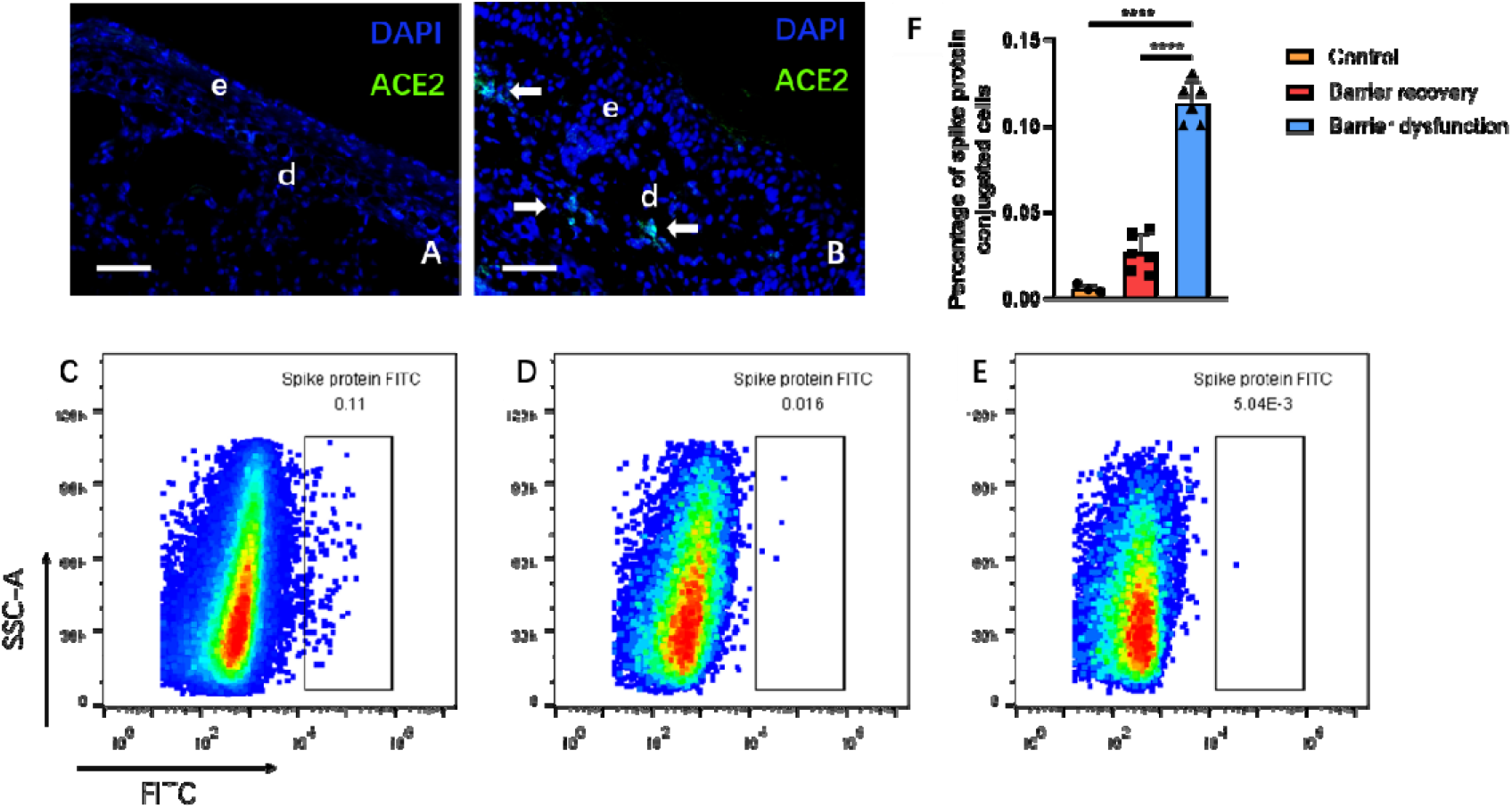
Deposition of FITC conjugated spike protein of mouse models with different skin condition. (A and B) Immunofluorescence stain of mouse without (A) or with (B) skin barrier dysfunction after applied with FITC conjugated spike protein. Bar indicated 10μm. Arrow indicated the deposition of spike protein. (C-E) Quantity of spike protein conjugated cell in the skin with barrier dysfunction and applied with FITC marked spike protein without (C) or with (D) barrier recovery moisture and the skin without barrier dysfunction (E) applied with FITC marked spike protein detected by FACS. (F) The comparison of the percentage of spike protein conjugated cells in the skin of different groups with skin barrier function. Bars indicate mean±SD of experiment groups. ** P < 0.05, *** P < 0.01 and **** P<0.001 P < 0.01.

### SARS-COV-2 pseudovirus could be detected in the dermal of mouse skin with inflammatory status and deteriorated skin barrier

Since the size of spike protein is much smaller than the actual SARS-COV-2, there is limitation of our data showed above. We further used SARS-COV-2 Pseudovirus-Luciferase (Sino Biological, PSV001), which contained recombinant pseudotyped lentiviral particles with SARS-CoV-2 spike protein to mimic SARS-CoV-2 cell infection (Nie et al., 2020) and humanized ACE2 mice (Shanghai Model Organism) to verify if topical contact with SARS-COV-2 could lead to infection of mice with barrier dysfunction. The mice were divided into 3 groups (n=3), mice with barrier dysfunction with topically treatment of pseudo virus, mice with barrier dysfunction without treatment of pseudo virus and mice without barrier dysfunction treated with pseudo virus. There were reports that SARS-COV-2 could be found in water (Randazzo et al., 2020) and salmon All the groups of mice were reared in one box. The skin, intestine and lung were obtained 48hrs after virus treatment. Then immunofluorescence was prepared to the obtained tissues. The GFP cells could be detected in the skin and lung of mice treated with pseudo virus and with barrier dysfunction (Figure 2). The GFP cells could not be detected in the mouse model treated with pseudo virus without barrier dysfunction as well as the mouse with barrier dysfunction not treated with pseudo virus.

**Fig. 2.**
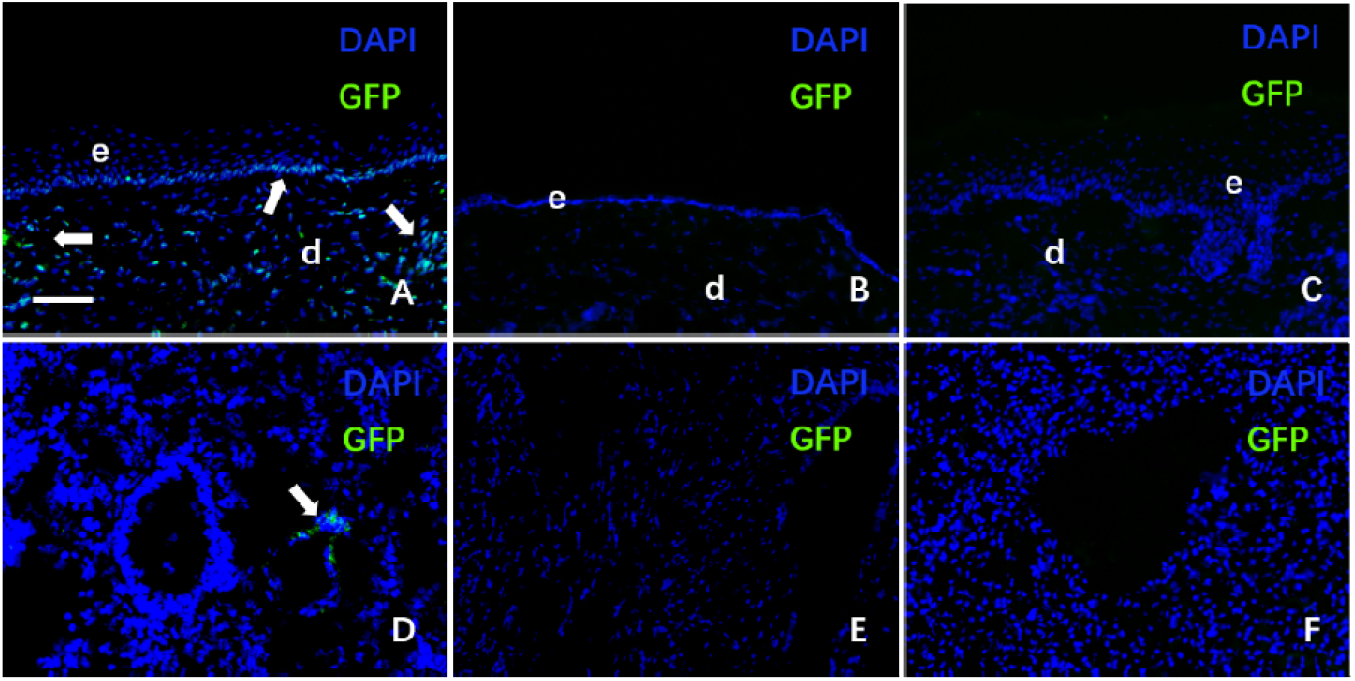
GFP cell expression in skin and lung of mouse models with or without barrier dysfunction as well as treated or untreated with SARS-COV-2 pseudo virus. (A)GFP cell expression in skin of mouse with barrier dysfunction and treated with SARS-COV-2 pseudo virus. Bar indicated 10μm (B)GFP cell expression in skin of mouse without barrier dysfunction and treated with SARS-COV-2 pseudo virus. (C) GFP cell expression in skin of mouse with barrier dysfunction without treatment of SARS-COV-2 pseudo virus. (D) GFP cell expression in lung of mouse with barrier dysfunction and treated with SARS-COV-2 pseudo virus. (F) GFP cell expression in lung of mouse without barrier dysfunction and treated with SARS-COV-2 pseudo virus. (G) GFP cell expression in lung of mouse with barrier dysfunction without treatment of SARS-COV-2 pseudo virus..

### Elevated ACE2 expression in the lesion of psoriasis and atopic dermatitis

To evaluate whether skin inflammatory disorders with barrier dysfunction have elevated ACE2 level which might enhance the possibility of transmitting of SARS-CoV-2 in skin, we randomly select 5 samples of psoriasis, 5 samples of atopic dermatitis which were all with skin barrier dysfunction and 3 samples without barrier dysfunction (All obtained from patients with nevus). Immunohistochemistry (IHS) stain was performed on all the samples. The ACE2 was expressed mainly in the stratum basale and dermis (Fig. 3, A to C). In the IHC stain of atopic dermatitis ACE2 positive stain could observed on the higher layer when compared with psoriasis and skin without barrier dysfunction. While in the dermis, the ACE2 positive cells in psoriasis and atopic dermatitis were recognized mostly as infiltrated inflammatory cells. The ACE2 expression in the IHC stain was measured by Image J IHC profiler plugin. ACE2 expression of psoriasis and atopic dermatitis is significantly elevated as compared with the skin without barrier dysfunction (Fig. 3D). These results support that skin ACE2 level of inflammatory skin disorders with barrier dysfunction were upregulated compared with the skin without barrier dysfunction.

**Fig. 3.**
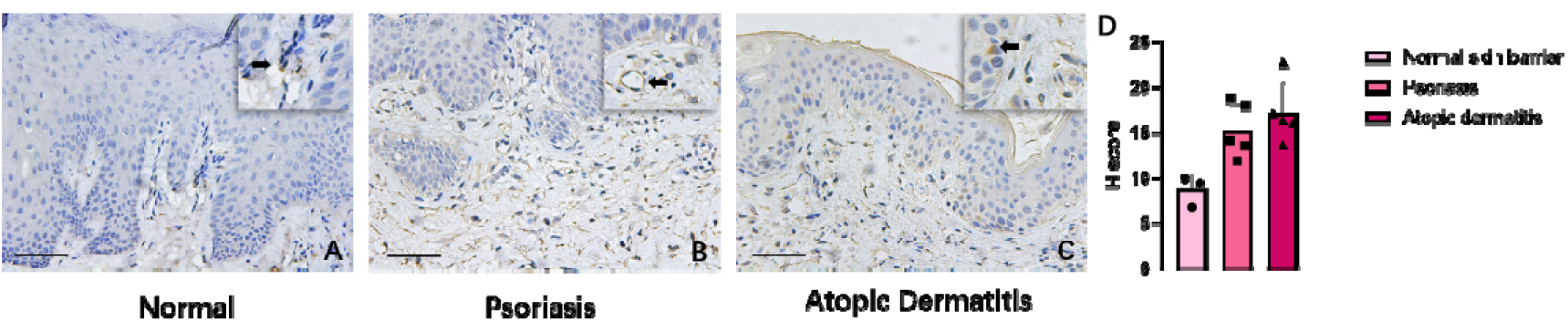
The IHC stain of ACE2 in the skin tissue with different skin conditions. (A-C) The ACE2 expression of skin with normal skin barrier function (A), with psoriasis (B) and with atopic dermatitis (C). Bar indicated 10μm. Arrow indicated positive stain. (D) The comparison of ACE2 expression in the skin with normal barrier function, lesion of psoriasis patient and lesion of patient with atopic dermatitis Bars indicate mean±SD of experiment groups ** P < 0.05 Student’s t test

**Fig. 4.**
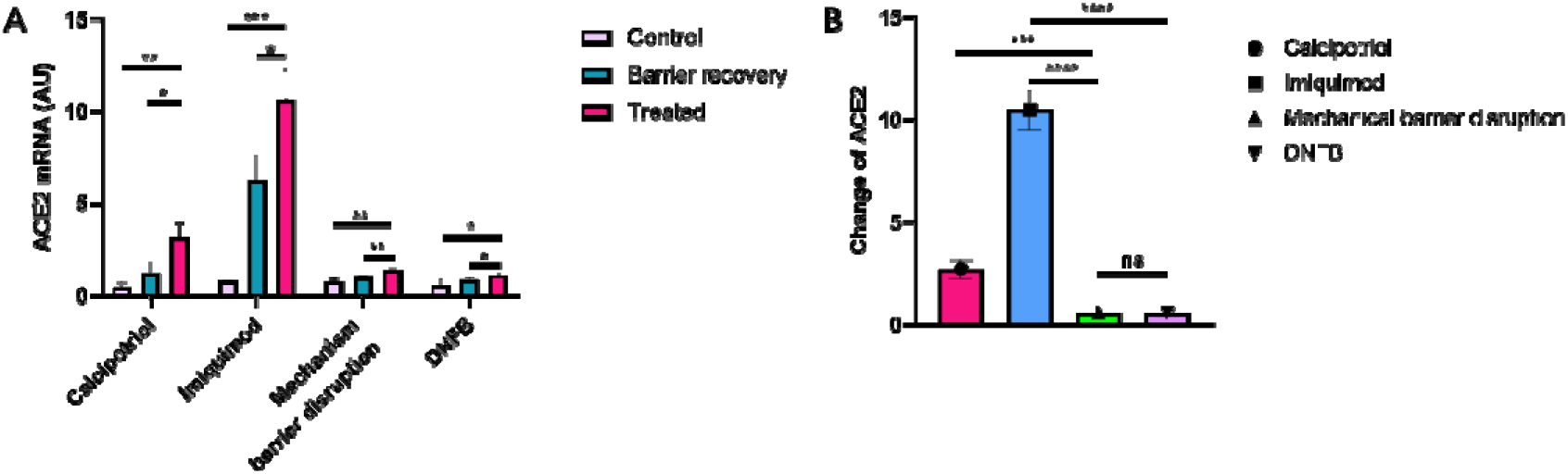
The ACE2 mRNA expression of different inflammatory disorders with skin barrier dysfunction mouse models. (A) Comparison of ACE2 mRNA expression of different inflammatory disorders with skin barrier dysfunction mouse models. (B) Comparison of ACE2 mRNA expression change of model establish before and after in different mouse model groups. Bars indicate mean±SD of experiment groups. ** P < 0.05, *** P < 0.01 and **** P<0.001 P < 0.01. Student’s t test.

### Elevated ACE2 expression in inflammatory skin disorders with barrier dysfunction mouse models and could be reduced by topical skin recovery moisture

Then does other inflammatory skin disorders with barrier dysfunction share the same higher ACE2 expression? We test this in four different mouse models, including psoriasis, atopic dermatitis mouse models induced by imiquimod or calcipotriol respectively, acute skin inflammatory skin disorder induced repeated tape-stripping(Wang et al., 2015) as well as 1-Fluoro-2,4-dinitrobenzene (DNFB). All four models were recognized as mouse models of inflammatory skin disorder with deteriorated barrier function. Paired control without skin inflammatory skin disorder and barrier dysfunction was established in the same length of time (Imiquimod control group was applied with vaseline while calcipotriol and DNFB group control was applied with absolute ethanol, mechanism skin barrier dysfunction control was established by eliminate mechanism treatment). The ACE2 mRNA expression of imiquimod group, calcipotriol group, mechanical barrier disruption group and DNFB group were all significant elevated compared with paired group control (Fig. 4A). Furthermore, the deviation of ACE2 mRNA expression in imiquimod induced psoriasis mouse model and calcipotriol induced atopic dermatitis mouse model were higher compared with mechanism disruption induced and DNFB induced mouse models with barrier dysfunction(Fig. 4B).

Besides the results above, when applied with skin recovery moisture (Linoleic acid-ceramide moisturizer recognized as sufficient in prevent psoriasis and atopic dermatitis relapse through skin barrier recovery and alleviate inflammation. Shanghai Jahwa Corporation) (Liu et al., 2015; Yang et al., 2019) the mRNA expression of ACE2 in the imiquimod treated group and calcipotriol treated group was downregulated as compared with the skin recovery moisture untreated group (Fig. 4a).

### IL-17 antibody could reduce the ACE2 expression in the lesion of psoriasis

To evaluate the influence of biologics on the inflammatory skin with barrier dysfunction, we randomly select 5 psoriasis patients who were under the treatment of IL-17 antibody (Taltz, Eli Lilly and Company). The lesional skin of patients was obtained on week 0 and week 8 after first use of the IL-17 antibody. ACE2 mRNA expression detection was prepared as previously described (Wang et al., 2014). The skin ACE2 mRNA expression of patients underwent using IL-17 antibody for 8 weeks is downregulated compared with the ACE2 expression in the skin on week 0 when the IL-17 antibody treatment started (Fig. 5A). To confirm the result, we randomly select 3 patients under the IL-17 antibody treatment and skin of patients were biopsied on week 0 and week 8 after first use of the IL-17 antibody. The tissue was prepared for immunofluorescence stain as previously described (Li et al., 2011). The immunofluorescence stain revealed the fluorescence intensity of ACE2 downregulated in the patient skin underwent IL-17 antibody treatment on week 8 compared with the patient skin before the IL-17 antibody (Fig. 5, B to D). Data above, either mRNA or protein of ACE2 from psoriasis patients, showed biologics might contribute to the defensing of SARS-CoV-2 in inflammatory skin disorders with barrier dysfunction through modulating skin barrier function, downregulating ACE2 expression and alleviate the inflammation.

**Fig. 5.**
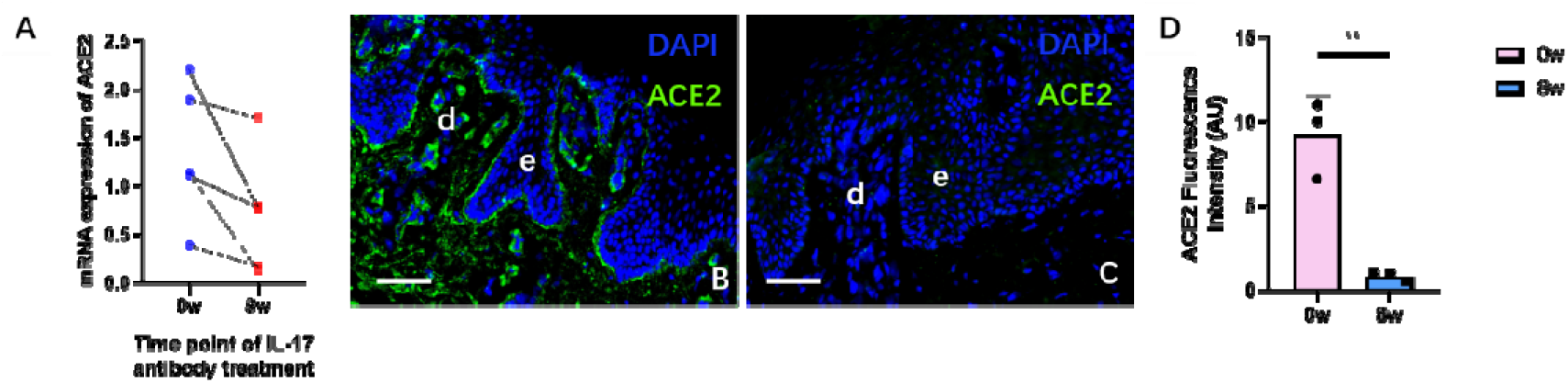
The expression of ACE2 in the lesion before and after the IL-17 antibody treatment. (A) The skin ACE2 mRNA expression in the lesion before (0w) and 8 weeks after (8w) the IL-17 antibody treatment. (B) The comparison of ACE2 immunofluorescence stain in the patient skin before and 8 weeks after the IL-17 antibody treatment. (C and D) Immunofluorescence stain of ACE2 in the patient skin that underwent IL-17 antibody treatment before (C) and 8 weeks after (D). Bar indicated 10μm. Bars indicate mean±SD of experiment groups. ** P < 0.05, Student’s t test.

## Discussion

With the pandemic of COVID-19, the method to prevent SARS-COV-2 from spreading become more and more urgent. Reveal the possible way of SARS-COV-2 transmitting may encourage new method reduce the COVID-19 spreading. The expression of ACE2 in skin gives the possibility that skin might be another link to the COVID-19 transmission. We use spike protein and pseudo virus showed that skin could be the potential target of SARS-COV-2 and the infection risk could be affected by the cutaneous conditions. Data recently published that ACE2 expressed in skin and in atopic dermatitis, a typical inflammatory skin disorder with barrier dysfunction, ACE2 gene level was elevated (Radzikowska et al., 2020), we added that besides atopic dermatitis, another inflammatory skin disorder with barrier dysfunction held elevated ACE2 level in lesion. There is one thing should be noted that the expression of ACE2 is not on the surface of skin, however, it mainly expressed in the dermal and the cells between dermal and epidermal. Deteriorated skin barrier gives various microbes the opportunity to get through the baseline of epidermal and so does SARS-COV-2 especially in the inflammation state. In our study we recognized that besides the already known physical dysfunction of the skin barrier which may lead the COVID-19 S protein binding the ACE2 expressing cell in the stratum basale and dermal, upregulation of ACE2 in the skin with barrier dysfunction were also detected in our study. These dermal ACE2 positive cells held an infiltrating inflammatory cell look. And there’s a possibility that these cells could be macrophage (Sluimer et al., 2008) or dendritic cells (Yang et al., 2020) which indicated that inflammatory atmosphere along with the deteriorated skin barrier could be the front door SARS-COV-2 knocked on. The elevated ACE2 level in the skin inflammatory disorder with deteriorated barrier provide the suitable environment for the transmission of SARS-COV-2. We also divided the skin inflammatory disorder with deteriorated barrier models into different groups and the models related more to inflammation (Bäsler and Brandner, 2017; Egawa and Kabashima, 2016; Jensen and Proksch, 2009) seemed to have higher ACE2 expression. As in our previous study the relieve of psoriasis was followed by the recovery of skin barrier, our data now revealed the skin ACE2 expression downregulated along with the alleviation of psoriatic skin barrier dysfunction due to IL-17 antibody treatment which could be a support that IL-17 antibody treatment in psoriasis might affect the COVID-19 transmission in a preventive way. Besides that, skin barrier recovery moisture which could also alleviated the inflammation as well as protect the skin barrier from dysfunction showed potential in reducing the risk of COVID-19 infection transmission. Deteriorated barrier function makes the virus easier to enter the skin, while the increased expression of ACE2 makes the virus easier to enter the cell. Inflammatory skin disorders (including psoriasis and atopic dermatitis) are accompanied by deteriorated barrier function and increased ACE2 expression, therefore, it is possible to increase the risk of COVID-19. (Fig. 5) Our work not only provided a new insight into the COVID-19 transmission way and it could be an alert to those whose skin exposure to the virus without a proper protection.

## STAR+METHODS

### Pseudo virus infecting mice

The imiquimod treatment was prepared as topical applied imiquimod each ear 10mg per day for 5 days. The pseudo virus (Sino Biological, cat PSV001) treatment was prepared as depositing the virus in PBS and made it 2×10^3^ copies/ml (Randazzo et al., 2020). Then 30ul virus was applied per mouse ear on day 6. The mice were divided into three groups as group A treated with imiquimod (3M) and virus, group B treated with vehicle cream (Vaseline Lanette cream, Fargon) as well as PBS, and group C treated with imiquimod and PBS. The mice were sacrificed 48 hours after applying the virus or PBS.

### Human tissue specimens

All procedures and use of (anonymized) tissue were performed according to recent national ethical guidelines. Human skin were obtained from patients undergoing biopsy procedures for diagnostic purposes or surgery for various reasons.

### Immunohistochemistry

Tissue samples were fixed in 10% buffered formalin, dehydrated, and embedded in paraffin. Paraffin sections, 6 μm thick, were stained with haematoxylin and eosin (H&E) to verify the diagnosis. ACE2 and Ki-67 IHC staining was performed on 4-μm-thick paraffin sections fixed on Superfrost Plus slides (Menzel Gläser, Braunschweig, Germany). Deparaffinization and antigen retrieval were performed in Target Retrieval Solution with pH 9 (ACE2) or pH 6 (Ki-67) at 97°C for 20 min using the PT Link platform (Dako, Glostrup, Denmark). Subsequently, the sections were washed in TBS (tris-buffered saline and incubated with primary antibodies directed against ACE2 (1:100, abcam, cat: ab15348) and Ki-67 (1:100, MIB-1, Dako) in an Autostainer Link48 automated staining platform (Dako) for 20 min at room temperature. The slides were then washed in TBS and visualization was performed using the EnVision FLEX system (Dako) according to the manufacturer’s instructions.

### Immunofluorescent staining

Frozen cryosections of skin from patients and mice with the indicated genotype or treatment were fixed in ice-cold acetone for 10 min before staining. The following antibodies were used for immunofluorescent staining, Goat Anti-Rabbit IgG H&L AlexaFluor 488 (abcam, Cat:ab150077)Rabbit polyclonal to Angiotensin Converting Enzyme 2 (abcam,Cat : ab15348). 4’,6-Diamidino-2-phenylindole (DAPI) (Fluka) was used for the staining of nuclei. Antibodies were diluted in antibody diluent (Dako). Immunostained samples were analyzed with a fluorescence microscope (Olympus DP72). Isotype IgGs were used as control stainings.

### Skin barrier dysfunction model

Mice (4-6 wks) were used in building the model. Psoriasis mouse model was established by topically treating each ear with 10 mg of 5% imiquimod cream (Imiquimod Cream 5%; 3M Pharmaceuticals) daily for 5 days. Atopic dermatitis mouse model were established by applying 3 nmol MC903 (MedChemExpress, Cat. No.: HY-10001) each ear per day for 10 days. Mechanical barrier disruption mouse model was achieved by stripping both sides of the earlobe with cellophane tape (3M Schotch) seven times. Contact dermatitis mouse model was established by applying 10ul 0.15% DNFB (Sigma-Aldrich, Inc., St. Louis, MOon the ear in day 0. 5 days later the ear was rechallenged by 10ul 0.15% DNFB and contact dermatitis like skin was induced.

### Quantitative RT-PCR

RNA extraction and PCR were carried out as follows. Primer sequences used were designed for murine ACE2 FORWARD: TGGGCAAACTCTATGCTGACTG, murine ACE2 BACKWARD: CCCTTCATTGGCTCCGTTTCT. Murine GAPDH FORWARD: CCTCGTCCCGTAGACAAAATG GAPDH BACKWARD: TGAGGTCAATGAAGGGGTCGT. Murine β-actin FORWARD: AGCCATGTACGTAGCCATCC β-actin BACKWARD: CTCTCAGCTGGTGGTGAA. Human ACE2 FORWARD: CATTGGAGCAAGTGTTGGATCTT, human ACE2 BACKWARD: GAGCTAATGCATGCCATTCTCA. Human GAPDH FORWARD: CACATGGCCTCCAAGGAGTAA HUMAN GAPDH BACKWARD: TGAGGGTCTCTCTTCCTCTTGT. Human CD209 FORWARD: GGATACAAGAGCTTAGCAGGGTG HUMAN CD209 BACKWARD: GCGTGAAGGAGAGGAGTTGC.

### Reproduction of FITC conjugated S-protein and protein applying method

Dissolve proteins and FITCs in FluoroTag™ FITC Conjugation Kit(Sigma-Aldrich)kit buffer. FITC was slowly added to the recombinant novel coronavirus spike glycoprotein (Cusabio, cat: CSB-EP3324GMY1) while stirring. Than the mixture was coverred with foil and stir at room temperature for 2 hours. After that FITC conjugated protein was collected through the G-25. FITC conjugated S-protein was than applied to the ear skin of mouse model with barrier dysfunction and group control. 24 hours later the treated part of skin was taken and underwent immunofluorescence observation.

### Skin barrier recovery

Mice were applied with skin barrier recovery moisture (Linoleic acid-ceramide moisturizer,Shanghai Jahwa Corporation) 5 days before processing the skin barrier deteriorating method. Skin barrier recovery moisture was applied 10mg each ear per day.

